# Genetic Variations and Altered Blood mRNA Level of Circadian Genes and BDNF as Risk Factors of Post-Stroke Cognitive Impairment among Eastern Indians

**DOI:** 10.1101/2023.07.07.548090

**Authors:** Dipanwita Sadhukhan, Arindam Biswas, Smriti Mishra, Daytee Maji, Parama Mitra, Priyanka Mukherjee, Gargi Podder, Biman Kanti Ray, Atanu Biswas, Tapas Kumar Banerjee, Subhra Prakash Hui, Ishani Deb

## Abstract

**Background:** Post-stroke cognitive impairment (PSCI) is a clinical outcome in around 30% of post-stroke survivors. BDNF is a major gene in this regard. It regulates and being regulated by circadian rhythm. The circadian genes are correlated with stroke timings at molecular level. However, studies suggesting the role of these on susceptibility to PSCI is limited.

**Aim:** We aim here to determine a) genetic risk variants in circadian clock genes, *BDNF* and b) dysregulation in expression level of *CLOCK*, BMAL1 and *BDNF*, that may be associated with PSCI.

**Methods:** BDNF (rs6265G/A, rs56164415C/T), CLOCK (rs1801260T/C, rs4580704G/C) and CRY2 (rs2292912C/G) genes variants were genotyped among 119 post-stroke survivors and 292 controls from Eastern part of India. In addition, we analysed their gene expression in PBMC from 15 PSCI cases and 12 controls. The mRNA data for BDNF was further validated by its plasma level through ELISA.

**Results:** Among the studied variants, only rs4580704/*CLOCK* showed an overall association with PSCI (P = 0.001) and lower BMSE score. Its ‘C’ allele showed a correlation with attention deficiency. The language and memory impairments showed association with rs6265/*BDNF* while the ‘CC’ genotype of rs2292912/*CRY2* negatively influenced language and executive function. A significant decrease in gene expression for *CLOCK* and *BDNF* in PBMC (influenced by specific genotypes) of PSCI patients was observed than controls. Unlike, Pro-BDNF, plasma level mBDNF was also lower in them.

**Conclusions:** Our results suggest that the circadian genes and *BDNF* play a role in PSCI on both genetic and transcript level.

## Introduction

Stroke is a leading cause of death and is considered as an important cause of long-term disability and cognitive impairment including dementia worldwide. Approximately 30% of stroke survivors go on to develop dementia [1]. Several cognitive domains mainly attention, memory, language, and orientation are get affected [2]. Owing to the complex nature of stroke, and inconstant post-stroke outcomes among individuals many studies attempted to identify ethnicity dependent genetic risk factors worldwide. With this regard, the primary gene of interest is *BDNF* and which is a fundamental molecule involved in neuronal survival, differentiation, and synapse plasticity – the cellular processes implicated in learning and memory. Former studies found cognitive impairment to be associated with the changes in peripheral BDNF levels [3]. Along with the *cis-*regulatory mechanism [4], there is also a master circadian control of BDNF expression and release involving the PGC1α/irisin/BDNF axis [5] and AMP/MAPK/CREB signalling pathway [6]. It is also reported that desynchronization or impairments in circadian rhythm often lead to the development of several disorders including cognitive loss like AD and progressive PD, etc. as evident from genetic association studies and *in-vivo* model systems resulting in the identification of associated variants and underlying mechanisms [7].

There are data derived from both humans and experimental animals indicating a direct connection between circadian clock genes and neuronal cell death following ischemic injury [8-10]. However, the role of the core clock gene towards susceptibility of post-stroke cognitive impairment was not evaluated properly. Since, BDNF is a prime factor related to cognition as well as one of the clock-controlled genes, we hypothesize that “*not only BDNF but circadian clock genes also may confer susceptibility towards post-stroke cognitive impairment and there may be a change in expression of circadian clock components in PSCI patients*”.

In this study, we aim to (1) evaluate functional/regulatory SNVs in *BDNF* and core clock genes *i*.*e BMAL1* and *CLOCK* that code for transcription factors as potent risk factors for post-stroke cognitive impairment; (2) correlate selected SNVs with different cognitive domains and characterize the mRNA expression of BDNF, CLOCK in peripheral blood mononuclear cells (PBMCs) and (3) estimate plasma mature BDNF (mBDNF) and Pro-BDNF level which may be correlated with poor cognitive outcome after stroke.

## Materials and Methods

### Patients and Controls

The present study was conducted upon the post stroke survivors recruited from the National Neurosciences Centre Calcutta and Bangur Institute of Neurosciences, IPGME&R, Kolkata. Informed consent taken as per guidelines of the Indian Council of Medical Research (ICMR). All the cohorts were older than 18 years with neuroimaging evidence of acute stroke. All had Bengali as their mother tongue, able to comprehend and speak and able to read and write. On the other hand, cases with dysphasia or any other form of language deficit, significant psychiatric illness, substance abuse, and history of psychotropic drug intake, drugs known to cause cognitive impairment and major head injury were excluded from the study. The demographic details of patients are summarised in Table 1. The history of known risk factors for stroke including hypertension, diabetes mellitus, hyperlipidemia, ischemic heart disease were obtained from the patients’ medical files while data on alcohol consumption or smoking were collected by asking verbal questions. In addition, age and ethnicity matched unrelated healthy controls with no personal or family history of stroke or any neurological symptoms were recruited in the present study from Kolkata.

**Table 1:** Demographic characteristics of Post-Stroke Cognitive Impaired and Post-Stroke Cognitive Normal groups.

| Demographic details / Subjects | PSCI (N = 46; 39%) | PSCN (N = 73; 61%) | p-value |
| --- | --- | --- | --- |
| Age range (years) | 28 - 78 | 28 - 79 |  |
| Age at onset ± SD | 55.04 ± 10.99 | 49.12 ± 10.25 | **0.0029** |
| Male | 34 (74%) | 63 (86%) | 0.092 |
| Female | 12 (26%) | 10 (14%) |  |
| Familial history of stroke | 22 (49%) | 41 (56%) | 0.455 |
| Hypertension | 35 (78%) | 58 (79%) | 0.821 |
| Diabetes | 20 (44%) | 22 (30%) | 0.165 |
| Previous known heart disease | 1 (2.22%) | 7 (9.59%) | 0.153 |
| Smoking/Alcohol | 20 (44%) | 45 (61.64%) | 0.087 |
| High blood cholesterol (>200 mg/dl) | 2 (4.44%) | 11 (15.07%) | 0.127 |
| High blood triglyceride (>150 mg/dl) | 2 (4.44%) | 9 (12.33%) | 0.202 |
| BMI | 23.09 ± 3.37 | 22.72 ± 2.82 | 0.766 |
| NIHSS (at admission) | 8.27 ± 4.19 | 7.36 ± 5.07 | 0.343 |
| MRS (at admission) | 2.41 ± 1.12 | 2.22 ± 1.23 | 0.511 |
| Recurrent event of stroke | 10 (22.22%) | 16 (21.92%) | 1.000 |
| Death after stroke | 2 (4.44%) | 1 (1.36%) | 0.557 |

PSCI: Post-Stroke Cognitive Impaired; PSCN: Post-Stroke Cognitive Normal

### Cognitive Parameters

Cognitive assessment was carried out within 6 to 12 months of stroke. All underwent comprehensive cognitive tests for attention, language, memory, visuospatial skill, and executive functions, etc. [11,12]. The different subdomain analysis for cognition was done in details [11].

### Selection of SNVs

On basis of a) contradictory reports on association of following SNVs like rs1801260C/T, rs4580704G/C of *CLOCK*, rs6265G/A of *BDNF*, rs2292912C/G of *CRY2* with neurological diseases in different populations; b) evidence of functional role of these SNVs from experimental data, eQTL (expression quantitative trait locus) findings reported in on the GTEx portal website (www.gtexportal.org) and rSNP base prediction (http://rsnp.psych.ac.cn/quickSearch) the present study aimed to genotype these variants for their potential association with the disease in our study cohort.

### Genomic DNA isolation and Genotyping of SNVs

Genomic DNA was isolated from fresh whole blood by a conventional salting out method using sodium perchlorate followed by isopropanol precipitation [13] and then dissolved in TE (10 mM Tris–HCl, 0.1 mM EDTA, pH 8.0). Polymerase chain reaction (PCR) was carried out using the specific primer pairs. Genotypes for SNVs were determined by restriction digestion of its PCR products with suitable enzymes (New England Biolabs Inc, Ipswich) following manufacturers’ protocol. The digested products were separated by 7% polyacrylamide gel electrophoresis, and visualized by ethidium bromide staining. About 10% of the samples for each variants were randomly selected for sequencing to rule out genotyping errors.

### Gene Expression Analysis

For gene expression study, peripheral blood mononuclear cells (PBMCs) were first isolated from fresh whole blood collected from stroke patients admitted in hospitals and from healthy controls using Histopaque 1077 (Sigma-Aldrich, St. Louis, US) double-gradient density centrifugation. Total RNA from PBMC was extracted using Trizol reagent (ThermoFisher Scientific, US) and dissolved in DEPC treated water, stored at -80^0^C. Afterwards, equal amount of RNA (2 μg) from each sample was subjected to cDNA synthesis using reverse transcriptase Kit (Promega Corporation, USA) following standard procedures. With these cDNAs the quantitative real time PCR (qPCR) was performed to measure *CLOCK, BMAL1 and BDNF* expression using gene specific primers (CLOCK_F: 5’-GGGGCAGTCATGGTACCTAG -3’; CLOCK_R: 5’-GCCTGAGATGGTTGCTGAAC-3’ BMAL1_F: 5’-TGAAGACAACGAACCAGACA -3’; BMAL1_R: 5’-CGTGCCGAGAAACATATTCC-3’ and BDNF_F: 5’-GTATTAGTGAGTGGGTAACGG -3’; BDNF_R: 5’-GCACTTGGTCTCGTAGAAGTA-3’) designed with the web-based software Primer3 and Syber Green PCR Master mix (Applied Biosystems, US). While, for normalisation purpose expression of *GAPDH* (Glyceraldehyde 3-phosphate dehydrogenase) (GAPDH_F: CTGACTTCAACAGCGACACC; GAPDH_R: TGCTGTAGCCAAATTCGTTGT) was considered as endogenous control. Here, the QuantStudio 5 Real-Time PCR (Applied Biosystems, US) instrument was programmed for quantitation of Ct values for each gene.

### Measurement of Plasma level of mBDNF and Pro-BDNF

The blood samples of the PSCI, Post-stroke cognitive normal (PSCN) were collected before 12 a.m. (to exclude interference from circadian rhythm). Fasting blood samples were centrifuged at 3,000×g for 20 mins after 60 min of incubation. Then isolated plasma was subjected to 12000×g for 15 min at 4^0^C for preparation of platelet poor plasma (PPP). For the measurement of mature BDNF (mBDNF) and Pro-BDNF using mature and Pro-BDNF Rapid ELISA Kit (Biosensis, Thebarton, Australia), the platelet free supernatant was diluted in assay diluent in 1:20 dilution ratio followed by addition to the precoated microplate wells and incubation for 45 min (mBDNF) and 2 hrs (Pro-BDNF) respectively. Then, after five washes, respective detection antibody into each well was added and incubated for 30 min. Again solution was discarded and washed as described above. Finally, the 1 × streptavidin–HRP conjugate into each well was added and incubated for 30 min followed by discarding the solution and washing as described above again. Finally, TMB and stop solution were added according to the instructions. Absorbance values for each sample were read at 450 nm on a plate reader (Varioskan Lux, Thermo Scientific).

### Statistical Analysis

Hardy-Weinberg equilibrium (HWE) at the polymorphic sites was tested using a chi square test with one degree of freedom. Genotype association was evaluated for p-value, odds ratio and 95% confidence interval (CI) using Javastat (http://statpages.info/ctab2x2.html). Appropriate corrections of significance values were also applied using the Benjamini-Hochberg correction method (false-discovery rate – FDR values) because of multiple SNVs were investigated. The q values of less than 0.05 were considered to be significant (https://tools.carbocation.com/FDR).

Statistical analysis for gene expression data was performed in Graph Pad Prism 5.0 using Mann-Whitney U-test to compare cases and controls, and Kruskal-Wallis test to compare the expression between different genotypes studied here. Both tests were two-tailed with a significance level p < 0.05. Data presented as mean log2 transformed expression ± SEM. Expression quantitative trait loci (e QTL) data was accessed from Genotype –Tissue Expression (GTEX project) website (www.gtexportal.org).

## Results

### Demographic characteristics of study subjects

One hundred and nineteen post-stroke survivors (mean age at onset: 51.53 ± 10.95, Male: Female = 4.41: 1) were included in the present study. Age and ethnicity matched unrelated healthy controls (n=292, mean age, 57.42 ± 9.35 years; Male: Female: 4.43:1) were collected. The demographic data and the principal variables for Ischemic stroke (IS) were compared between post-stroke survivors (PSCI vs PSCN) as shown in Table 1. Except the older age none of the traditional risk factors was found to be differently associated with cognitive impairment (P = 0.0029).

### Association of *BDNF (rs6265*G/A, *rs56164415*C/T*), CLOCK (rs1801260*T/C, *rs4580704*G/C*) and CRY2 (rs2292912*C/G) SNVs with Post stroke cognitive Impairment

A total of five different genetic variants in *CLOCK, BDNF, CRY2* were genotyped in 119 post -stroke survivors and 292 controls. The genotypic distribution of studied SNVs were within Hardy-Weinberg equilibrium and no linkage disequilibrium (LD) between the variants on same gene. Table 2 shows the allele and genotype frequency of these SNVs. The ‘C’-allele [OR = 2.502; 95% CI = 1.498 - 4.179; P-value = 0.001], genotypes with C allele [OR = 3.226; 95% CI = 1.627-6.398; P-value = 0.001] of rs4580704 were observed to be associated with risk for PSCI (35%) among eastern Indians when compared to healthy controls (17.47%) and PSCN individuals (25%). However, in the cases of rs6265, rs56464415 of *BDNF*, rs1801260 of *CLOCK* and rs2292912 of *CRY2* neither allelic nor genotypic association were found with the disease.

**Table 2:** Distribution of genotype and allele frequencies of *BDNF, CLOCK* and *CRY2* SNPs among Stroke patients and Controls.

| **rs6265 (BDNF)** |  |  |  |  |  |  |  |
| --- | --- | --- | --- | --- | --- | --- | --- |
| **Group** | **Genotype Frequency (%)** |  |  | **Allele Frequency (%)** |  | **p-value** | **Odds Ratio (95%CI)** |
|  | **GG** | **GA** | **AA** | **G** | **A** |  |  |
| PSCI (n=46) | 20 (43.5%) | 23 (50%) | 3 (6.5%) | 63 (68.5%) | 29 (31.5%) | 0.558 | G vs A: 0.820 (0.463 – 1.451) |
| PSCN (n=73) | 41 (56%) | 24 (33%) | 8 (11%) | 106 (72.6%) | 40 (27.4%) | 0.193 | GG vs GA+AA: 0.600 (0.285 – 1.263) |
| Controls (n = 280) | 132 (47.14%) | 138 (49.28%) | 10 (3.58%) | 402 (71.78%) | 158 (28.21%) | *0.535 | *G vs A: 0.854 (0.530 – 1.375) |
|  |  |  |  |  |  | *0.750 | *GG vs GA+AA: 0.862 (0.460 – 1.617) |
| **rs56164415 (BDNF)** |  |  |  |  |  |  |  |
|  | **CC** | **CT** | **TT** | **C** | **T** | **p-value** | **Odds Ratio (95%CI)** |
| PSCI (n=44) | 23 (52%) | 20 (45.5%) | 1 (2.5%) | 66 (75%) | 22 (25%) | 0.449 | C vs T: 1.313 (0.718 – 2.400) |
| PSCN (n=69) | 33 (48%) | 30 (43.5%) | 6 (8.5%) | 96 (69.5%) | 42(30.5%) | 0.702 | CC vs CT+TT: 1.195 (0.560 – 2.547) |
| Controls (n = 280) | 155 (55.3%) | 103 (36.7%) | 22 (8%) | 413 (73.7%) | 147 (26.3%) | *0.896 | *C vs T: 1.068 (0.636 – 1.792) |
|  |  |  |  |  |  | *0.746 | *CC vs CT+TT: 0.883 (0.467 – 1.670) |
| **rs1801260 (CLOCK)** |  |  |  |  |  |  |  |
|  | **TT** | **TC** | **CC** | **T** | **C** | **p-value** | **Odds Ratio (95%CI)** |
| PSCI (n=44) | 18 (41%) | 23 (52%) | 3 (7%) | 59 (67%) | 29 (33%) | 0.325 | T vs C: 1.348 (0.770 – 2.360) |
| PSCN (n=69) | 22 (32%) | 39 (57%) | 8 (11%) | 83 (60%) | 55 (40%) | 0.420 | TT vs TC+CC: 1.479 (0.674 – 3.245) |
| Controls (n = 292) | 118 (40.41%) | 144 (49.31%) | 30 (10.28%) | 380 (65%) | 204 (35%) | *0.810 | *T vs C: 1.092 (0.679 – 1.758) |
|  |  |  |  |  |  | *1.000 | *TT vs TC+CC: 1.021 (0.536 – 1.945) |
| **rs4580704 (CLOCK)** |  |  |  |  |  |  |  |
|  | **GG** | **GC** | **CC** | **G** | **C** | **p-value** | **Odds Ratio (95%CI)** |
| PSCI (n=39) | 16 (41%) | 19 (49%) | 4 (10%) | 51 (65%) | 27 (35%) | 0.201 | C vs G: 1.554 (0.836 – 2.887) |
| PSCN (61) | 36 (59%) | 19 (31%) | 6 (10%) | 91 (75%) | 31 (25%) | 0.102 | GC+CC vs GG: 2.070 (0.914 – 4.686) |
| Controls (n = 292) | 202 (69.18%) | 78 (26.71) | 12 (4.11 %) | 482 (82.53%) | 102 (17.47%) | *0.001 | *C vs G: **2.502 (1.498 – 4.179)** |
|  |  |  |  |  |  | *0.001 | *GC+CC vs GG: **3.226 (1.627 – 6.398)** |
| **rs2292912 (CRY2)** |  |  |  |  |  |  |  |
|  | **CC** | **GC** | **GG** | **C** | **G** | **p-value** | **Odds Ratio (95%CI)** |
| PSCI (n=39) | 15 (38.46%) | 22 (56.41%) | 2 (5.13%) | 52 (67%) | 26 (33%) | 0.232 | C vs G: 1.471 (0.810 – 2.668) |
| PSCN (n=59) | 14 (23.73%) | 40 (67.80%) | 5 (8.47%) | 68 (58%) | 50 (42%) | 0.174 | CC vs GC+GG: 2.009 (0.833 – 4.847) |
| Controls (n =276) | 104 (37.7%) | 152 (55.1%) | 20 (7.2%) | 360 (65%) | 192 (35%) | *0.899 | *C vs G: 1.067 (0.645 – 1.763) |
|  |  |  |  |  |  | *1.000 | *CC vs GC+GG: 1.034 (0.519 – 2.060) |

P value: PSCI vs PSCN; \*P value: PSCI vs Controls

Because multiple genetic variants were examined in our study cohort, further appropriate corrections of P values were made using the Benjamini-Hochberg correction method (false-discovery rate – FDR values). The significance of allelic association between rs4580704 and PSCI did withstand multiple testing (q value = 0.004, which is the adjusted p value for 4 variants) even after doing FDR.

Haplotypes were next determined based on the genotypes at rs6265 and rs56464415 of *BDNF* for PSCI and PSCN individuals using Haploview (version 4.2) (data not shown). However, we failed to find any disease associated protective or risk haplotypes between the groups.

### Association of *BDNF* and Circadian SNVs with Cognitive raw scores

A total of 119 right handed IS patients out of 198 stroke survivors were followed up within 3-12 months of onset for evaluation of cognitive status as described earlier [11] and 39% of cases had BMSE score ≤ 24. The “C” allele of rs4580704 of *CLOCK* and “CC” genotype of rs2292912 of *CRY2* showed an association with lower BMSE score referring an overall post-stroke cognitive impairment [Table 3].

**Table 3:** Comparison of Post-Stroke cognitive outcome based on genetic variants in circadian genes and BDNF.

|  | Attention | Language | Memory | Visuospatial | Executive | BMSE (<24) |
| --- | --- | --- | --- | --- | --- | --- |
| **rs6265 (BDNF)** | 10.50±4.93 | 33.37±7.38 | 35.21±11.63 | 5.83±2.98 | 8.82±6.11 | 24.37±5.80 |
| GG (n = 61) | P: 41 (67%)<br>I: 20 (33%) | P: 56 (92%)<br>I: 5 (8%) | P: 39 (64%)<br>I: 22 (36%) | P: 28 (46%)<br>I: 33 (54%) | P: 33 (54%)<br>I: 28 (46%) | P: 41 (67%)<br>I: 20 (33%) |
| GA+AA (n = 58) | 9.77±4.77<br>P: 37 (64%)<br>I: 21 (36%) | 29.40±10.02<br>P: 44 (76%)<br>I: 14 (24%) | 29.98±14.37<br>P: 20 (35%)<br>I: 38 (65%) | 5.69±3.26<br>P: 28 (48%)<br>I: 30 (52%) | 7.72±5.57<br>P: 27 (47%)<br>I: 31 (53%) | 23.72±5.63<br>P: 32 (55%)<br>I: 26 (45%) |
| P-value | 0.469<br>0.705 | **0.039**<br>**0.024** | **0.002**<br>**0.003** | 0.938<br>0.855 | 0.344<br>0.465 | 0.265<br>0.193 |
| **rs56164415 (BDNF)** | 9.91±4.96 | 31.48±9.46 | 32.31±13.90 | 5.80±3.09 | 8.14±6.25 | 23.57±6.30 |
| CC (n = 56) | P: 35 (62.5%)<br>I: 21 (37.5%) | P: 46 (82%)<br>I: 10 (18%) | P: 30 (54%)<br>I: 26 (46%) | P: 27 (48%)<br>I: 29 (52%) | P: 30 (54%)<br>I: 26 (46%) | P: 33 (59%)<br>I: 23 (41%) |
| CT+TT (n = 57) | 10.39±4.78<br>P: 39 (68%)<br>I: 18 (32%) | 31.49±8.72<br>P: 48 (84%)<br>I: 9 (16%) | 34.38 ±12.49<br>P: 31 (54%)<br>I: 26 (46%) | 5.61±3.17<br>P: 25 (44%)<br>I: 32 (56%) | 8.47±5.52<br>P: 26 (46%)<br>I: 31 (54%) | 24.36±5.28<br>P: 36 (63%)<br>I: 21 (37%) |
| P-value | 0.638<br>0.556 | 0.786<br>0.806 | 0.589<br>1.000 | 0.705<br>0.707 | 0.695<br>0.454 | 0.716<br>0.702 |
| **rs1801260 (CLOCK)** | 9.92±4.92 | 32.51±7.44 | 32.92±12.59 | 5.10±3.01 | 8.53±5.84 | 22.63±6.90 |
| TT (n = 40) | P: 32 (80%)<br>I: 16 (20%) | P: 36 (90%)<br>I: 4 (10%) | P: 21 (53%)<br>I: 19 (47%) | P: 13 (32.5%)<br>I: 27 (67.5%) | P: 20 (50%)<br>I: 20 (50%) | P: 22 (55%)<br>I: 18 (45%) |
| TC+CC (n = 73) | 10.33±5.00<br>P: 54 (74%)<br>I: 19 (26%) | 31.05±9.77<br>P: 58 (79%)<br>I: 15 (21%) | 33.71±13.58<br>P: 40 (55%)<br>I: 33 (45%) | 6.08±3.14<br>P: 41 (56%)<br>I: 32 (44%) | 8.26±6.07<br>P: 36 (49%)<br>I: 37 (51%) | 24.75±5.00<br>P: 47 (64%)<br>I: 26 (36%) |
| P-value | 0.423 | 0.809 | 0.709 | 0.083 | 0.759 | 0.176 |
|  | 0.417 | 0.193 | 0.846 | **0.019** | 1.000 | 0.420 |
| **rs4580704 (CLOCK)** | 11.07±4.81 | 31.70±7.97 | 33.05±12.95 | 5.96±2.95 | 8.63±5.13 | 24.52±5.37 |
| GG (n = 52) | P: 37 (71%) | P: 43 (83%) | P: 28 (54%) | P: 24 (46%) | P: 28 (54%) | P: 39 (75%) |
|  | I: 15 (29%) | I: 9 (17%) | I: 24 (46%) | I: 28 (54%) | I: 24 (46%) | I: 13 (25%) |
| GC+CC (n= 48) | 9.10±5.06 | 31.48±10.02 | 32.82±13.35 | 5.32±3.31 | 8.08±6.62 | 22.87±6.49 |
|  | P: 29 (60%) | P: 41 (85%) | P: 24 (50%) | P: 20 (42%) | P: 22 (46%) | P: 25 (52%) |
|  | I: 19 (40%) | I: 7 (15%) | I: 24 (50%) | I: 28 (58%) | I: 26 (54%) | I: 23 (48%) |
| P-value | **0.057** | 0.702 | 0.865 | 0.400 | 0.609 | 0.283 |
|  | 0.295 | 0.789 | 0.841 | 0.690 | 0.548 | **0.022** |
| **rs2292912 (CRY2)** | 9.49±5.32 | 30.64±11.32 | 30.64±11.32 | 5.609±3.44 | 5.71±3.06 | 21.31±6.82 |
| CC (n = 29) | P: 17 (59%) | P: 20 (69%) | P: 12 (41%) | P: 12 (41%) | P: 16 (55%) | P: 14 (48%) |
|  | I: 12 (41%) | I: 9 (31%) | I: 17 (59%) | I: 17 (59%) | I: 13 (45) | I: 15 (52%) |
| GC+GG (n = 69) | 10.35±4.93 | 32.86±6.10 | 35.10±11.42 | 5.71±3.06 | 8.48±6.17 | 24.64±5.37 |
|  | P: 47 (68%) | P: 62 (90%) | P: 38 (55%) | P: 32 (46.4%) | P: 34 (49%) | P: 45 (65%) |
|  | I: 22 (32%) | I: 7 (10%) | I: 31 (45%) | I: 37 (53.6%) | I: 35 (51%) | I: 24 (35%) |
| P-value | 0.468 | 0.887 | 0.069 | 0.744 | **0.020** | **0.022** |
|  | 0.486 | **0.016** | 0.270 | 0.665 | 0.661 | 0.174 |

Further, a detail analysis of cognitive domains into subdomains (attention, memory, language, etc.) followed by analyses of raw scores, identified ‘A’ allele rs6265/*BDNF* as genetic risk factor for language (P = 0.039, 0.024) and memory impairment (P = 0.002 and 0.003). In a similar way, language and executive functional ability was found to be negatively influenced by “CC” genotype of rs2292912/*CRY2* (P = 0.016; 0.020) while visuospatial impairment is more common among cases with “TT”genotype for rs1801260/*CLOCK* (P = 0.019). On the other hand, for attention domain, a trend was observed for “C” allele carrying genotypes of rs4580704/CLOCK with P value = 0.057.

### Gene expression analysis

Then, on the basis of existing knowledge about the role of circadian genes and *BDNF* in cognition especially learning and memory, a transcript analysis was performed in PBMCs from PSCI patients (N = 15) and controls (N = 12). The relative expression levels of *BDNF* and *CLOCK* was found to be significantly lower among cases with P = 0.0190, 0.008 respectively (for PSCI patients: mean log2 CLOCK expression = -2.492± .1022; mean log2 BDNF expression = -2.853 ± 0.1118; for Controls: mean log2 CLOCK expression = -1.894 ± .08455; mean log2 BDNF expression = -2.272 ± 0.1047) (Figure 1A, 1B). However, no such difference was observed for BMAL1 gene (Figure 1C).

**Legend to Figure 1:**
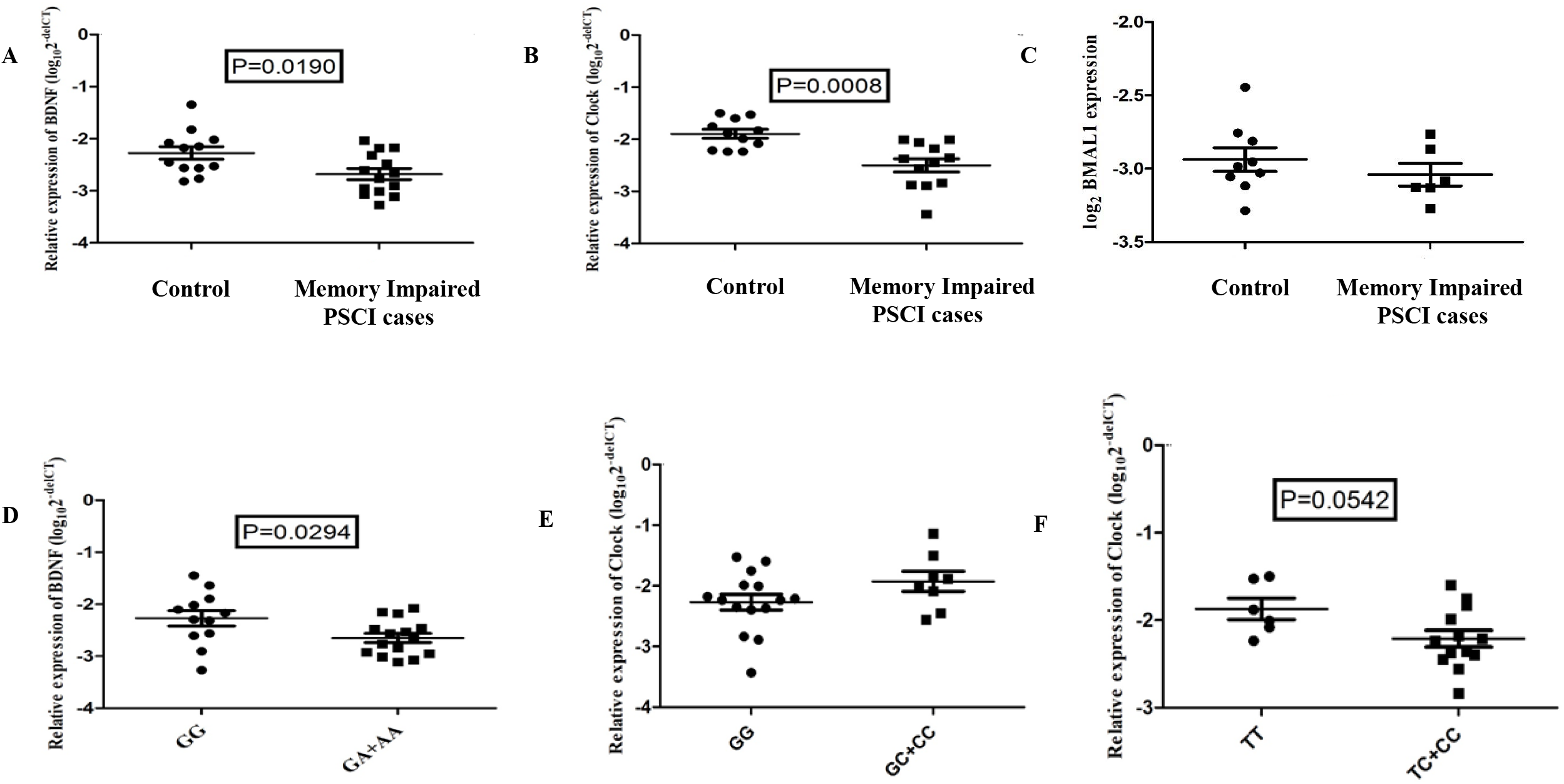
*BDNF, CLOCK, BMAL1* mRNA expression in correlation to disease status/genotype. **(A)** Quantification of BDNF mRNA levels in human PBMC from controls (n = 15) and Post stroke cognitive impaired (PSCI) patients (n = 12), normalized to the housekeeping gene GAPDH using Mann-Whitney U-test (P = 0.0190) ; **(B)** Quantification of CLOCK mRNA levels in same individuals (P = 0.0008); **(C)** Quantification of BMAL1 mRNA levels in same individuals; **(D)** BDNF mRNA expression levels among study subjects grouped by rs6265G/A genotypes; **(E)** comparison of log2 transformed CLOCK mRNA expression levels among individuals grouped by rs4580704G/C and **(F)** rs1801260 T/C genotypes. Data presented as mean ± standard deviation in all graphs. The significance level was set at P ≤ 0.05.

### CLOCK, BDNF mRNA expression in correlation to genotype

Next, to investigate whether *BDNF* and *CLOCK* gene expression can be linked to rs6265G/A, rs4580704G/C and rs1801260T/C variants, irrespective of disease phenotype the minor allele carrying genotypes were grouped together. By doing this, we found that mean gene expression is lower for the ‘A’ allele of rs6265/*BDNF* (−2.650 ± 0.08807) and ‘C’ allele of rs1801260/*CLOCK* (−2.211 ± 0.09572) between the groups (Figure 1D, F). Although a trend for higher expression was observed for ‘C’ allele of rs4580704/*CLOCK* (−1.928 ± 0.1648) but fails to meet significance (Figure 1E).

### Comparison of Plasma level of mBDNF level between PSCI and PSCN

Finally, with the aim to validate our expression data as well to identify differential conversion of Pro-BDNF into mBDNF (if any), the plasma level of both the forms were measured by ELISA among the study groups. Our biochemical analyses showed that plasma level of mBDNF (memory impaired PSCI= 4504 ± 534 pg/ml; PSCN = 6884 ± 672 pg/ml; t= 2.211; df =37, P = 0.0337) but not the Pro-BDNF (PSCI= 1358 ± 139 pg/ml; PSCN = 1243 ± 92.6 pg/ml; t= 0.7133; df =28, P = 0.4816) was significantly lower in PSCI group than PSCN individuals (Figure 2A). Plasma levels of mature BDNF also showed a significant positive correlation with the lower memory function score among PSCI group (P = 0.0232) (Figure 2B).

**Legend to Figure 2:**
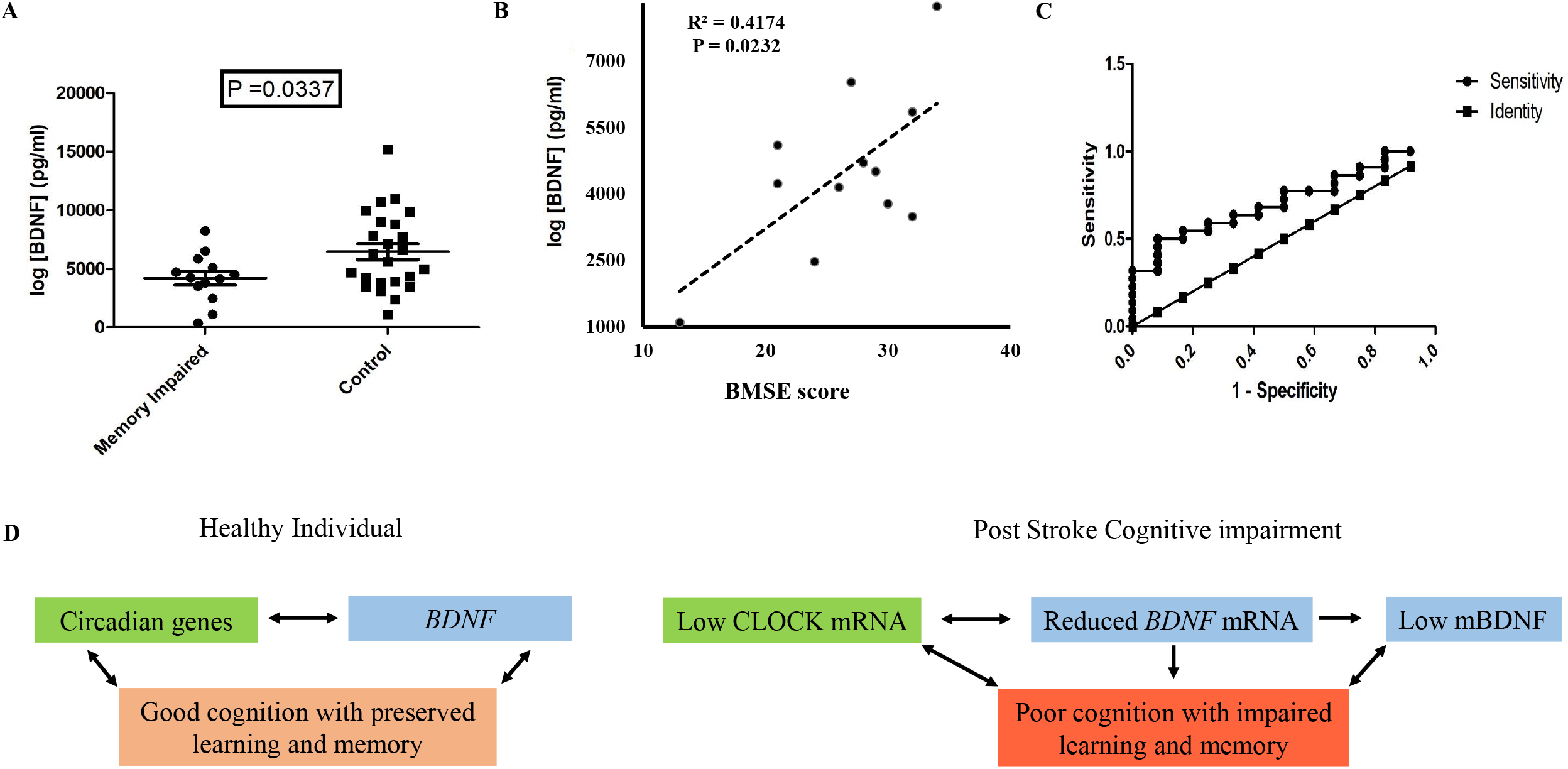
Change in plasma mature BDNF level. (A) P < 0.05 comparison between cases and controls; B) significant positive correlation between BMSE score and plasma mBDNF; C) ROC curve for plasma mBDNF level as diagnostic marker for post stroke memory impairment; D) Diagramatic representation showing dysregulation in *CLOCK* and *BDNF* level in PSCI.

ROC analysis was performed to compare the predictive value of plasma mBDNF for diagnosing PSCI. The results showed that plasma mBDNF had an AUC of 0.7159 ± 0.08778 (95% CI: 0.5438-0.8880; P = 0.04001). The optimal cut off for plasma mBDNF was 4595 ng/ml with 0.68 sensitivity and 0.58 specificity (Figure 2C).

## Discussion

In the present study, we have investigated the relationship of Circadian genes and *BDNF* with post stroke cognitive impairment among our Eastern Indian population. Of the major socio-demographic parameters, only increasing age showed a higher prevalence in cognitive impaired patients than cognitive preserved survivors. Our data is consistent with other world populations suggesting age as an independent risk factor for post-stroke cognitive decline [14,15]. On the other hand, *BDNF* being the primary gene responsible for plasticity, its association with post-stroke outcome has been tested in different population [16]. Although the Val66Met substitution of *BDNF* was found to alter activity-dependent release of BDNF, which consequently impacts the efficiency of BDNF-TrkB [17] signalling, its association with debility and neurological shortfall is ethnicity dependent [18-24]. In our study, GA + AA genotype/BDNF groups showed poor memory function and language difficulty in comparison to GG genotype. However, comparing other sub-domains, no difference in raw scores were found between two groups which is consistent with previous studies by Razei et al., 2017 [24] and Han et al., 2020 [3]. It is reported that the restoration of BDNF expression improves vascular cognitive impairment [25] in mice, while in human study subjects, differential BDNF level (mostly from serum and few from PBMC) during acute phase of stroke serves as predictor of functional outcome [26]. Given this background and owing to the transcriptional and post translation complexity [27] in formation of mature BDNF, here we examined both mRNA and protein level of BDNF among memory impaired PSCI cases and PSCN ones. PBMCs are known to migrate and infiltrate the ischemic brain tissue as an inflammatory response and show considerable overlap in gene expression with brain tissues [28]. Here, we detected significantly lower a) relative expression of *BDNF* gene in PBMC and b) mBDNF level in plasma without a change in Pro-BDNF level, among PSCI patients compared to control group which corroborates with our genetic data. An additional negative influence of ‘A’ allele on BDNF expression was observed in these individuals irrespective of disease phenotype. Therefore, we can assume that the positive contribution of other genes and environmental factors on cognition is responsible for the better cognitive performance among individuals with ‘A’ allele at rs6265/BDNF. Our findings on plasma BDNF level (for both mature and pro) and its correlation with memory function score is consistent with a previous study by Hassan TM et al,.2018 [29]. Therefore, from this data we can suggest that, alteration in BDNF transcript level but not the cleavage of its pro-peptide may be responsible for the reduced BDNF level in peripheral blood sample.

BDNF being a clock controlled gene [30] and involved in regulation of circadian rhythm [31], in the present study we moreover investigated the role of *CLOCK* and *CRY2* on PSCI susceptibility. Along with an overrepresentation of ‘C’ allele among global cognitive status impaired cases, a novel association of it with lower attention score was observed in our study subjects. On the other hand, higher frequency of ‘CC’ genotype of rs2292912/*CRY2* was found to be associated with language and memory problems. However, the reduced sample size after stratification and skewness of scores may have been prime factors for failing to meet the simultaneous significance level *i*.*e*. scoring as well frequency distribution for these circadian SNVs. These genetic variants previously reported to be associated with several disorders in different world populations [32]. However, for the first time, we have suggested these as potent risk factors for post-stroke cognitive outcome. Furthermore, in PBMC of memory impaired post stroke human subjects, a significantly lower relative expression level for *CLOCK* but not the *BMAL1* gene was detected when compared to controls. Although, an earlier study, in rat hippocampus identified a strong correlation between the daily expression of BDNF and Bmal1 and Cry1[33]. Unlike, our genetic data, here no difference in mRNA level between genotypes of rs4580704/CLOCK was observed. In contrast, for the reported 3’UTR variant, i.e rs1801260 [34], an effect of ‘C’ allele carrying genotype on decreased mRNA level was observed. According to eQTL (expression quantitative trait locus) data on the GTEx portal website (www.gtexportal.org) ‘CC’ genotype of rs4580704 / *CLOCK* and ‘GG’ genotype of rs2292912 (*CRY2*) correspond to respective increased and decreased level of mRNA expression in whole blood of healthy individuals. But no such overall findings in our case may be explained by disease status and involvement of other genetic and epigenetic modifiers [35]. Unfortunately, due to technical limitations, here we have failed to quantify and correlate the CRY2 mRNA level with disease.

In conclusion, our study first demonstrates that the genetic variants in *CLOCK, CRY2* serve as the susceptibility factors for PSCI. Furthermore, a differential expression analysis revealed alteration in the expression of *CLOCK* and *BDNF* genes may be implicated in the post stroke cognitive decline [Figure 2D]. Future researches on overall circadian rhythmicity and other clock controlled genes (ccgs) in PSCI is warranted to highlight it as potent therapeutic target.

## Acknowledgements

The authors are thankful to the patients, family members and healthy individuals who participated in the study.

## Author Contribution

Dipanwita Sadhukhan, Ishani Deb and Arindam Biswas were responsible for the concept, study design, experimental work and manuscript preparation. Smriti Mishra, Parama Mitra, Priyanka Mukherjee and Gargi Podder had put effort in data analysis (both clinical and genetic) and manuscript preparation. Atanu Biswas, Biman Kanti Ray, Tapas Kumar Banerjee evaluated patients and blood collections as being clinical collaborates. While, Subhra Prakash Hui provided the instrumentation facility in S. N. Pradhan Centre for Neurosciences, University of Calcutta and provide his input for manuscript preparation. All authors read the draft, provided their inputs and agreed on the final version of the manuscript.

## Data availability statement

The data that support the findings of this study are available from the corresponding author upon request.

## Ethics approval

All procedures performed in studies involving human participants were in accordance with the ethical standards of the institutional and national research committee as well as the 1964 Helsinki Declaration and its later amendments.

## Informed consent

Informed consent from all the participants were received prior to clinical data and sample collection

## References

1. Cullen B, O’Neill B, Evans JJ et al (2007) A review of screening tests for cognitive impairment. J Neurol Neurosurg Psychiatry 78:790–799. doi: 10.1136/jnnp.2006.095414

2. Al-Qazzaz NK, Ali SH, Ahmad SA et al (2014) Cognitive impairment and memory dysfunction after a stroke diagnosis: A post-stroke memory assessment. Neuropsychiatr Dis Treat 10:1677–1691. doi: 10.2147/NDT.S67184

3. Han Z, Qi L, Xu Q et al (2020) BDNF met allele is associated with lower cognitive function in poststroke rehabilitation. Neurorehabil Neural Repair 34:247–259. doi: 10.1177/1545968320902127.

4. Zheng F, Zhou X, Moon C et al (2012) Regulation of brain-derived neurotrophic factor expression in neurons. Int J Physiol Pathophysiol Pharmacol 4:188–200.

5. Zsuga J, More CE, Erdei T et al (2018) Blind spot for sedentarism: Redefining the diseasome of physical inactivity in view of circadian system and the irisin/bdnf axis. Front Neurol 9:818. doi: 10.3389/fneur.2018.00818.

6. Eckel-Mahan KL, Storm DR (2009). Circadian rhythms and memory: Not so simple as cogs and gears. EMBO Rep 10:584–59. doi: 10.1038/embor.2009.123.

7. Carter B, Justin HS, Gulick D et al (2021) The molecular clock and neurodegenerative disease: A stressful time. Front Mol Biosci 8:644–747. doi: 10.3389/fmolb.2021.644747.

8. Wiebking N, Maronde E, Rami A (2013) Increased neuronal injury in clock gene per-1 deficient-mice after cerebral ischemia. Curr Neurovasc Res 10:112–125. doi: 10.2174/1567202611310020004.

9. Tischkau SA, Cohen JA, Stark JT et al (2007) Time-of-day affects expression of hippocampal markers for ischemic damage induced by global ischemia. Exp Neurol 208:314–322. doi: 10.1016/j.expneurol.2007.09.003.

10. Beker MC, Caglayan B, Yalcin E et al (2018) Time-of-day dependent neuronal injury after ischemic stroke: Implication of circadian clock transcriptional factor bmal1 and survival kinase akt. Mol Neurobiol 55:2565–2576. doi: 10.1007/s12035-017-0524-4.

11. Biswas A, Sadhukhan D, Biswas A et al (2021) Identification of GBA mutations among neurodegenerative disease patients from eastern india. Neurosci Lett 751:135816. doi: 10.1016/j.neulet.2021.135816.

12. Das G, Dubey S, Sinharoy U et al (2021) Clinical and radiological profile of posterior cortical atrophy and comparison with a group of typical alzheimer disease and amnestic mild cognitive impairment. Acta Neurol Belg 121:1009–1018. doi: 10.1007/s13760-020-01547-4.

13. Johns MB, J., Paulus-Thomas JE (1989) Purification of human genomic DNA from whole blood using sodium perchlorate in place of phenol. Anal Biochem 180:276–278. doi: 10.1016/0003-2697(89)90430-2.

14. Srithumsuk W, Kabayama M, Gondo Y (2020) The importance of stroke as a risk factor of cognitive decline in community dwelling older and oldest peoples: the SONIC study. BMC Geriatr 20:24. https://doi.org/10.1186/s12877-020-1423-5.

15. Bao MH, Zhu SZ, Gao XZ, et al (2018) Meta-Analysis on the Association between Brain-Derived Neurotrophic Factor Polymorphism rs6265 and Ischemic Stroke, Poststroke Depression. J Stroke Cerebrovasc Dis 27:1599–1608. doi: 10.1016/j.jstrokecerebrovasdis.2018.01.010.

16. Levine DA, Wadley VG, Langa KM et al (2018) Risk Factors for Poststroke Cognitive Decline: The REGARDS Study (Reasons for Geographic and Racial Differences in Stroke). Stroke 49:987–994. doi: 10.1161/STROKEAHA.117.018529.

17. Chen ZY, Patel PD, Sant G et al (2004) Variant brain-derived neurotrophic factor (bdnf) (met66) alters the intracellular trafficking and activity-dependent secretion of wild-type bdnf in neurosecretory cells and cortical neurons. J Neurosci 24:4401–4411. doi: 10.1523/JNEUROSCI.0348-04.2004.

18. Di Lazzaro V, Pellegrino G, Di Pino G et al (2015) Val66met bdnf gene polymorphism influences human motor cortex plasticity in acute stroke. Brain Stimul 8:92–96.

19. Balkaya M, Cho S (2019) Genetics of stroke recovery: Bdnf val66met polymorphism in stroke recovery and its interaction with aging. Neurobiol Dis 126:36–46. doi: 10.1016/j.brs.2014.08.006.

20. Zhao J, Wu H, Zheng L et al (2013) Brain-derived neurotrophic factor g196a polymorphism predicts 90-day outcome of ischemic stroke in chinese: A novel finding. Brain Res 1537:312–318. doi: 10.1016/j.brainres.2013.08.061.

21. Kim JM, Stewart R, Park MS et al (2012) Associations of bdnf genotype and promoter methylation with acute and long-term stroke outcomes in an east asian cohort. PLoS One 7:e51280. doi: 10.1371/journal.pone.0051280.

22. Mirowska-Guzel D, Gromadzka G, Czlonkowski A et al (2012) Bdnf -270 c>t polymorphisms might be associated with stroke type and bdnf -196 g>a corresponds to early neurological deficit in hemorrhagic stroke. J Neuroimmunol 249:71–75. doi: 10.1016/j.jneuroim.2012.04.011.

23. Stanne TM, Tjarnlund-Wolf A, Olsson S et al (2014) Genetic variation at the bdnf locus: Evidence for association with long-term outcome after ischemic stroke. PLoS One 9:e114156. doi: 10.1371/journal.pone.0114156.

24. Rezaei S, Asgari-Mobarake K, Keshavarz P et al (2017) Brain-Derived Neurotrophic Factor (BDNF) Val66met (rs6265) Polymorphism Associated with Global and Multi-Domain Cognitive Impairment in Ischemic Stroke Patients. Activitas Nervosa Superior 59:28–36.

25. Niu Y, Wan C, Zhou B et al (2018) Aerobic exercise relieved vascular cognitive impairment via nf-kappab/mir-503/bdnf pathway. Am J Transl Res 10:753–761.

26. Karantali E, Kazis D, Papavasileiou V et al (2021) Serum bdnf levels in acute stroke: A systematic review and meta-analysis. Medicina (Kaunas)57. doi: 10.3390/medicina57030297.

27. Notaras M, & van den Buuse M (2019) Brain-derived neurotrophic factor (BDNF): Novel insights into regulation and genetic variation. The Neuroscientist 25:434–454.

28. Moore DF, Li H, Jeffries N et al (2005) Using peripheral blood mononuclear cells to determine a gene expression profile of acute ischemic stroke: a pilot investigation. Circulation 111:212–21. doi: 10.1161/01.CIR.0000152105.79665.C6.

29. Hassan TM, Yarube IU (2018) Peripheral brain-derived neurotrophic factor is reduced in stroke survivors with cognitive impairment. Pathophysiology 25:405–410. doi: 10.1016/j.pathophys.2018.08.003.

30. Liang FQ, Walline R, Earnest DJ (1998) Circadian rhythm of brain-derived neurotrophic factor in the rat suprachiasmatic nucleus. Neurosci Lett 242:89–92. doi: 10.1016/s0304-3940(98)00062-7.

31. D’Agostino Y, Frigato E, Noviello TMR et al (2022) Loss of circadian rhythmicity in bdnf knockout zebrafish larvae. iScience 25:104054. doi: 10.1016/j.isci.2022.104054.

32. Valenzuela FJ, Vera J, Venegas C et al (2016) Evidences of Polymorphism Associated with Circadian System and Risk of Pathologies: A Review of the Literature. Int J Endocrinol 2016:2746909. doi: 10.1155/2016/2746909.

33. Pramong R, Govitrapong P, Phansuwan-Pujito P (2017) Relationship between Circadian Clock Genes and the Neurotrophic Factor Genes in Rat Hippocampus. J Med Assoc Thai 100:S32–s39.

34. Ozburn AR, Purohit K, Parekh PK et al (2016) Functional implications of the clock 3111t/c single-nucleotide polymorphism. Front Psychiatry 7:67. doi: 10.3389/fpsyt.2016.00067.

35. Monti P, Iodice S, Tarantini L et al (2021) Effects of PM Exposure on the Methylation of Clock Genes in a Population of Subjects with Overweight or Obesity. Int J Environ Res Public Health 18:1122. doi: 10.3390/ijerph18031122.

